# The evolution of RNA interference among Metazoa

**DOI:** 10.1101/2023.05.08.538551

**Authors:** Alessandro Formaggioni, Gianmarco Cavalli, Mayuko Hamada, Tatsuya Sakamoto, Federico Plazzi, Marco Passamonti

## Abstract

In animals, three main RNA interference mechanisms have been described so far, which respectively maturate three types of small noncoding RNAs (sncRNAs): miRNAs, piRNAs and endo-siRNAs. The diversification of these mechanisms is deeply linked with the evolution of the Argonaute gene superfamily since each type of sncRNA is loaded by a specific Argonaute homolog protein. Moreover, other protein families play pivotal roles in the maturation of sncRNAs, like the DICER ribonuclease family, whose DICER1 and DICER2 paralogs maturate respectively miRNAs and endo-siRNAs. Among Metazoa, the distribution of these families has been only studied in major groups, and there are very few data for clades like Lophotrochozoa. Thus, we here inferred the evolutionary history of the animal Argonaute and DICER families including 43 lophotrochozoan species. Phylogenetic analyses along with newly sequenced sncRNA libraries depicted a loss of the endo-siRNA pathway along the Lophotrochozoa evolution, with the absence of DICER2 in Nematoda and Polyzoa, and with the absence of DICER2 and the Argonaute homolog in the rest of Trochozoa phyla. On the contrary, early diverging phyla, Platyhelminthes and Syndermata, showed a complete endo-siRNA pathway. On the other hand, miRNAs were revealed the most conserved and ubiquitous mechanism of the metazoan RNA interference machinery, confirming their pivotal role in animal cell regulation.

## Introduction

Argonaute proteins are cytoplasmic proteins that play a key role in most of the RNA interference (RNAi) pathways. They interact with a small non-coding RNA (sncRNA), forming an RNA-induced silencing complex (RISC). This ribonucleoprotein complex binds and silences target transcripts, using the complementary sncRNA as a probe (Iwakawa and Tomari 2022).Argonaute proteins can be found throughout most eukaryotic clades and share a common structure, featuring four domains: N-terminal (N), PIWI-Argonaute-Zwille (PAZ), Middle (MID) and P element-induced wimpy testis (PIWI) (Kuhn and Joshua-Tor 2013). The PIWI domain resembles the RNAse H domain’s structure, but only some Argonautes have been reported to cleave the target mRNA (Song et al. 2004). All the other Argonaute proteins repress the target trough proteins that interact with the RISC complex (Huntzinger and Izaurralde 2011; Wu et al. 2020).

Among the Argonaute superfamily, four different families have been characterized: *Trypanosoma*-AGO family, WAGO family, AGO-like family, and PIWI-like family (Swarts et al. 2014). *Trypanosoma*-AGO proteins have been identified only in the order Trypanosomatida (Garcia Silva et al. 2010). Conversely, WAGOs have been characterized in nematodes only, whereas AGO-like and PIWI-like proteins seem to be found in all animal phyla (Höck and Meister 2008; Swarts et al. 2014).

Three classes of metazoan interfering sncRNAs can be identified: micro-RNAs (miRNAs), small interfering RNAs (siRNAs), and piwi-interacting RNAs (piRNAs) (Iwakawa and Tomari 2022). Each class is matured by different pathways, is loaded by a different Argonaute protein and plays different cellular functions. Precursors of miRNAs (i.e., pri-miRNAs) are encoded by specific genes, transcribed by the RNA polymerase II (Y. Lee et al. 2004; Bartel 2018). Pri-miRNAs feature a hairpin secondary structure, with single-stranded ends at their 3’ and 5’ (Bartel 2018). These precursors undergo several maturation steps, starting immediately inside the nucleus, where they are targeted by the Microprocessor complex, which cleaves the overhanging nucleotides of pri-miRNAs, leaving a stem-loop structure with a 2bp offset, named pre-miRNA (Lee et al. 2003). Then, pre-miRNAs are exported in the cytoplasm, where they become substrate for DICER1, an endonuclease featuring one PAZ domain and two RNase III domains (Bernstein et al. 2001; Y.S. Lee et al. 2004). Through its catalytic activity, DICER1 removes the hairpin’s loop, leaving a ∼22bp ds-miRNA (Bernstein et al. 2001; Bartel 2018). Eventually, an AGO-like Argonaute binds to the ds-miRNA in the cytoplasm and disposes of one of the two strands, consequently resulting in a RISC complex. The miRNA-guided RISCs mostly target mRNAs by binding their 3’-UTR, interfering with their stability (Bartel 2018).

Unlike miRNAs, siRNAs may apparently be obtained from roughly every RNA capable of assuming a dsRNA structure (Shabalina and Koonin 2008). Therefore, siRNAs can originate from transcripts of transposons, repeated elements, or pseudogenes (Czech et al. 2008; Tam et al. 2008; Malone and Hannon 2009). Although some siRNAs have exogenous dsRNA precursors (e.g., viral RNAs), most of them have an endogenous origin, thus they are known as endo-siRNAs (Shabalina and Koonin 2008; Gammon and Mello 2015). Endo-siRNAs undergo a maturation pathway that is very similar to that of miRNAs, leading to the hypothesis that they evolved from a common ancestral RNAi system (Shabalina and Koonin 2008; Moran et al. 2017). Once the dsRNA precursors reach the cytoplasm, they are processed by DICER2, a paralog of DICER1, but with a DExH helicase domain instead of a PAZ domain. DICER2 cleaves a ∼21bp dsRNA that is loaded into an AGO-like Argonaute, called AGO2 in fruit flies (Y.S. Lee et al. 2004; Matranga and Zamore 2007; Czech et al. 2008). The resulting siRISC complex maintains one of the two strands, again using it as a probe (Matranga and Zamore 2007). The siRNA-mediated RNAi activity is not restricted to mRNAs, as siRNAs are known to partake in transposon silencing, DNA methylation, and antiviral defense (Galiana-Arnoux et al. 2006; Malone and Hannon 2009; Gammon and Mello 2015). A singular siRNA amplification pathway has been reported in nematodes, whose genomes encode RdRP enzymes, capable of using cleaved targets of the siRISC as a template to synthesize secondary siRNAs (Matranga and Zamore 2007).

Finally, piRNAs are 24-35 nt-long RNAs that originate from longer single strand precursors, consisting of either active transposons or transcripts of genomic piRNA clusters (Hirakata and Siomi 2016). Once these ssRNA precursors are exported through the nuclear pores, they undergo a rather complex maturation pathway, which takes place mostly in the perinuclear nuage (Weick and Miska 2014; Hirakata and Siomi 2016). Although piRNAs appear to be Metazoa-restricted (Grimson et al. 2008), biogenesis pathways vary significantly between clades (Weick and Miska 2014). Mature piRNAs interact with PIWI-like Argonautes, resulting in a piRISC that operates as a defense system against transposons, by means of its nuclease activity (Malone and Hannon 2009). Moreover, piRISCs have been linked to specific epigenetic modifications of the chromatin (i.e. H3K9me3) (Le Thomas et al. 2013). A peculiar piRNA amplification pathway called “ping-pong cycle” has been characterized in fruit flies, and later discovered in mice as well (Brennecke et al. 2007; Weick and Miska 2014). This amplification pattern is performed by two different PIWI-like Argonautes, namely one that binds sense-piRNAs (AGO3 in fruit flies) and one that loads antisense ones (Aubergine in fruit flies). As soon as one of these two proteins cleaves its target, it yields a secondary piRNA that can be loaded on the other PIWI-like protein, generating an amplification loop (Weick and Miska 2014; Hirakata and Siomi 2016). Another type of piRNA amplification is called “phasing”. PIWI loads a single strand RNA (pre-pre-piRNA) from the 5’ end, directing the endonucleolytic cleavage of the ribonuclease Zucchini at the 3’ end. This process is repeated along the pre-pre-piRNA leading to phased matured piRNAs (Mohn et al. 2015; Ozata et al. 2019).

The evolution and diversification of the Argonuate superfamily is deeply linked with the emergence of new RNAi pathways since each type of sncRNA is loaded by a different Argonaute homolog. miRNAs, piRNAs and endo-siRNAs were likely to be in the last metazoan common ancestor, since they have been described in Porifera, Cnidaria and most of the metazoan phyla (Grimson et al. 2008; Wheeler et al. 2009; Moran et al. 2013; Praher et al. 2017; Calcino et al. 2018; Fridrich et al. 2020). In some clades piRNA and endo-siRNA pathways have been secondarily lost (Wynant et al. 2017; Fontenla et al. 2021), while the miRNA pathway is likely to be ubiquitous in Metazoa (Fromm et al. 2022). 6

RNAi mechanisms are by far less studied in Lophotrochozoa compared to Deuterostomia and Ecdysozoa. This superclade is comprised by several phyla, such as Platyhelminthes, Syndermata, Annelida, and Mollusca, showing an astonishing variability. For medical and nutritional reasons, most of the data are restricted to Mollusca and Platyhelminthes. Parasitic Platyhelminthes (i.e., Neodermata) lacks the PIWI pathway, but the miRNA and endo-siRNA pathways have been reported (Fontenla et al. 2017; Fontenla et al. 2021). The PIWI pathway has been confirmed in Mollusca, where it is visible a clear signature of ping-pong amplification (Jehn et al. 2018). Moreover, many miRNAs have been annotated in mollusks (Fromm et al. 2022). Outside those phyla, few data have been published. The endo-siRNA pathway seems absent in the Annelida *Capitella teleta* (Khanal et al. 2022) and the annotation of miRNA family is restricted to one Syndermata, one Brachiopoda and two Annelida species (Fromm et al. 2022).

During last years, the increase of -omics data have made possible to compare and study the evolution of protein families along Lophotrochozoa. In this study, we exploited various -omics resources from nine lophotrochozoan phyla to annotate and characterize the diversification of the Argonaute and DICER proteins. We also analyzed sncRNA libraries to annotate the three sncRNA types and confirm the presence or absence of a particular sncRNA type in some phyla. The pattern reported by our results has never been described in any eukaryotic clade. Along the Lophotrochozoa evolution the endo-siRNA pathway has been progressively lost, starting with DICER2 in all Trochozoa (i.e., Nemertea, Phoronida, Brachiopoda, Annelida and Mollusca) and Polyzoa (Entoprocta + Bryozoa), followed by the loss of the fruit fly AGO2-like proteins. This pattern is confirmed by the distribution of DICER2 and AGO2-like proteins in the analyzed organisms. On the contrary, the piRNA and miRNA pathways appeared to be conserved in almost all Lophotrochozoa.

## Results

We annotated Argonaute proteins of 43 lophotrochozoan species analyzing 19 proteomes, 16 genomes and 8 transcriptomes. Argonaute sequences of *Homo sapiens* (Chordata), *Drosophila melanogaster* (Arthropoda), *Caenorhabditis elegans* (Nematoda) and *Nematostella vectensis* (Cnidaria) were retrieved from SwissProt and included in the phylogenetic analysis as references. Moreover, we include metazoan species whose RNAi pathways have been already studied (see Introduction), testing whether our Argonaute and DICER annotation matches results from literature. We chose proteomes of *Danio rerio* and *Gallus gallus* (Chordata), *Asterias rubens* (Echinodermata), *Anopheles gambiae* (Arthropoda), *Strongyloides ratti* (Nematoda) and the *Acropora muricata* (Cnidaria) genome assembly.

All the annotated Argonaute proteins were aligned, and the Maximum-Likelihood (ML) tree was inferred. The PIWI and AGO proteins of *Trypanosoma brucei* were obtained from Uniprot and used as outgroups. The phylogenetic tree of the Argonaute superfamily supported the known main families (Fig. 1; supplementary figure S1). The WAGO family, which is restricted to nematodes, included WAGOs and CSR-1 sequences from *C. elegans*. The PIWI-like family was characterized by AGO3, PIWI and AUB of *D. melanogaster*, HILI and HIWI of *H. sapiens* and PRG-1 of *C. elegans;* the AGO-like family comprised AGO1,2 of *D. melanogaster*, AGO1-4 of *H. sapiens* and ALG1,2 of *C. elegans*. Every family was widely supported by UFBoot and the SH-alrt test (Fig. 1).

**Figure 1.**
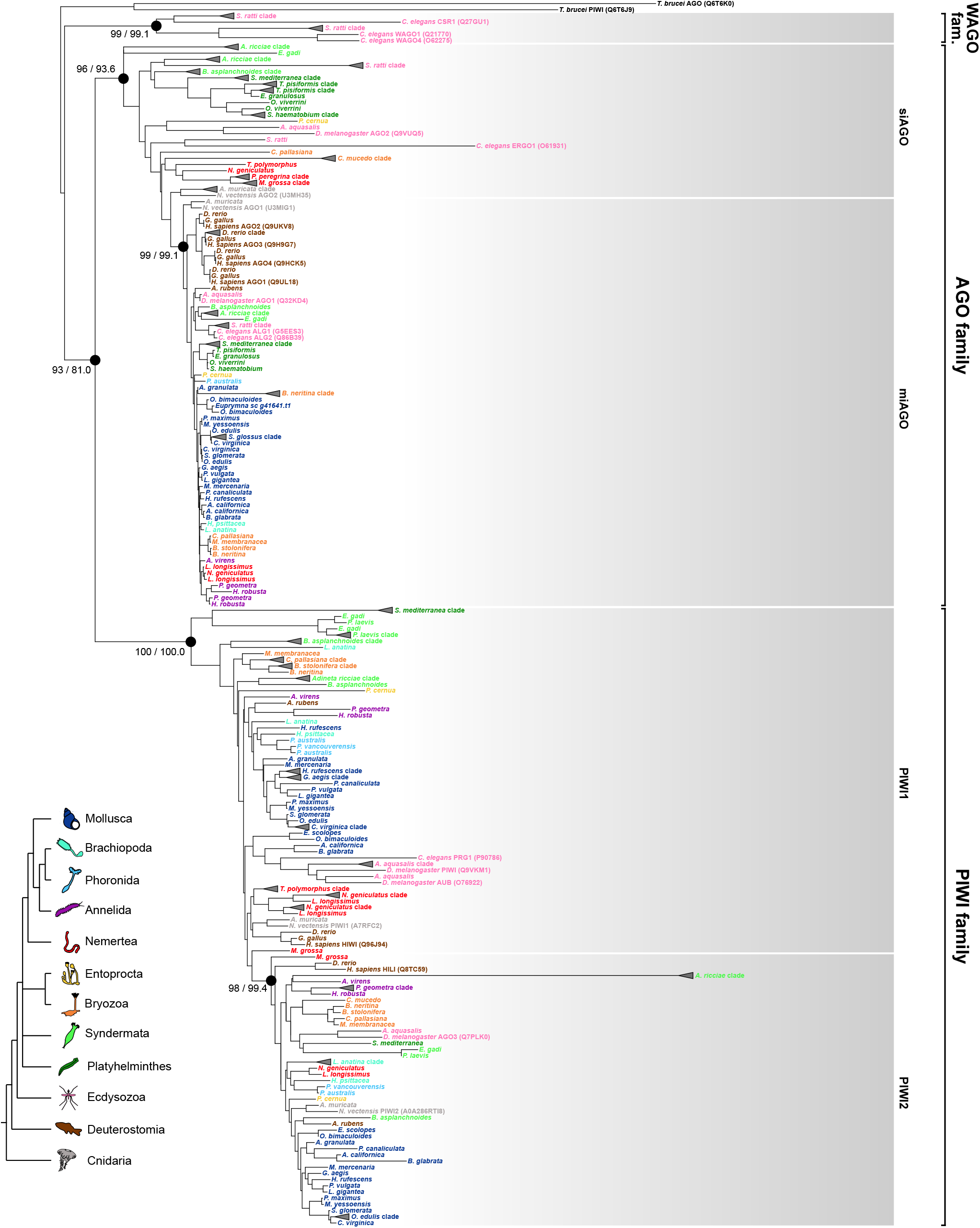
ML tree of the lophotrochozoan Argonaute proteins. For the six marked nodes the label shows UFBoot/SH-alrt value. Clades formed by paralogs of the same species were collapsed and represented with a triangle. For reference proteins the Uniprot accession code is reported in brackets. Species are colored according to the phylum. The color legend on the bottom left reconstructs the main phylogenetic relationships according to the ML-tree of Kocot and colleagues (2016).

The AGO-like family was divided in two groups. One monophyletic group (UFBoot = 99 and SH-alrt = 99.1) included *H. sapiens* AGO1-4, *D. melanogaster* and *N. vectensis* AGO1 and *C. elegans* ALG1,2: all these proteins interact with miRNAs. Almost all species featured a protein clustering within this clade, with very few exceptions, and at least one organism from every phylum was recovered in this clade (Fig. 1; supplementary table S1). We will refer to this clade as the “miAGO clade”. The second group within the AGO-like family was polyphyletic. *D. melanogaster* AGO2, *N. vectensis* AGO2 and *C. elegans* ERGO1 clustered within this clade. All these proteins target endo-siRNAs, thus we will henceforth call this group the “siAGO group”. Few lophotrochozoan phyla are included in this group, since we annotated at least one siAGO protein for all the Platyhelminthes and Entoprocta species, while three other phyla featured at least one species within the group, namely Nemertea, Bryozoa and Syndermata. We did not retrieve any siAGO protein from the remaining clades, namely Mollusca, Annelida, Brachiopoda, Phoronida (Fig. 1; supplementary table S1).

Similarly to the AGO-like family, protein clustering in the PIWI-like family can be divided in two groups: one is monophyletic and one is polyphyletic. The monophyletic group (UFBoot = 98 and SH-alrt = 99.4) comprised the *H. sapiens* HILI, the *N. vectensis* PIWI2 and the *D. melanogaster* AGO3; the polyphyletic group included *H. sapiens* HIWI, the *N. vectensis* PIWI1, the *C. elegans* PRG1 and the *D. melanogaster* AUB and PIWI. Each phylum was present in both groups with at least one species (supplementary table S1). PIWI-like proteins were almost absent in Platyhelminthes, apart from *Schmitdea mediterranea* (Fig. 1).

Regarding the six non-lophotrochozoan species included in the analysis, the annotation of Argonautes proteins matched the expectations: miAGO proteins were retrieved for all six species; siAGO proteins were absent in *D. rerio, G. gallus* and *A. rubens* (Deuterostomia); all six species reported PIWI1 and PIWI2 proteins, exception made for *S. ratti* (i.e. that lacked of both PIWI proteins) and, unexpectedly, *G. gallus* that lacked of PIWI2 (Fig. 1; supplementary table S1).

The phylogenetic analysis highlighted the presence of miAGO proteins in each phylum, but only some of them feature siAGO proteins. In Arthropoda, the preliminary structures of siRNAs and miRNAs are processed by two distinct DICER paralogs: DICER2 and DICER1, respectively (Shabalina and Koonin 2008). DICER2 has been found in other phyla, like Cnidaria and Platyhelminthes (Mukherjee et al. 2013a), while clades lacking siAGO proteins lack DICER2 as well. We annotated DICER proteins and inferred the phylogeny to understand whether Lophotrochozoa follow the same pattern.

We annotated DICER proteins querying the lophotrochozoan sequences against an annotated metazoan DICER set (Mukherjee et al. 2013a) and looking for the ribonucleases 3 domain. The phylogenetic analysis included the metazoan DICER set of Mukherjee and colleagues (2013) as references, including the *Zea mays* DICER proteins as outgroups (Fig. 2; supplementary figure S2). The resulting ML tree clustered the DICER proteins in two distinct groups. Recall the position of the reference sequences, DICER1 and DICER2 sequences are accordingly split in the two groups, apart from *Litopenaeus vannamei* DICER2, which is basal to all the other proteins. The monophyly of the DICER1 group is supported by the SH-alrt test (86.4) and the UFBoot (96). On the contrary, the low support values of the DICER2 node (UFBoot= 58 and SH-alrt = 64.3) undermine the hypothesis of a unique common origin of DICER2 proteins. Nonetheless, the presence of DICER2 was restricted to few lophotrochozoan phyla: all Platyhelminthes and three Syndermata species showed a DICER2 protein. On the other hand, DICER1 was annotated for every phylum: thus, Mollusca, Annelida, Phoronida, Brachiopoda, Nemertea, Bryozoa and Entoprocta reported DICER1 proteins, but no DICER2 proteins (Fig. 2; supplementary table S1). In line with the expectations, DICER2 was reported for *A. muricata, A. gambiae*, while it was lacking in *D. rerio, G. gallus, A. rubens* and *S. ratti*. On the contrary, DICER1 was annotated in all of them.

**Figure 2.**
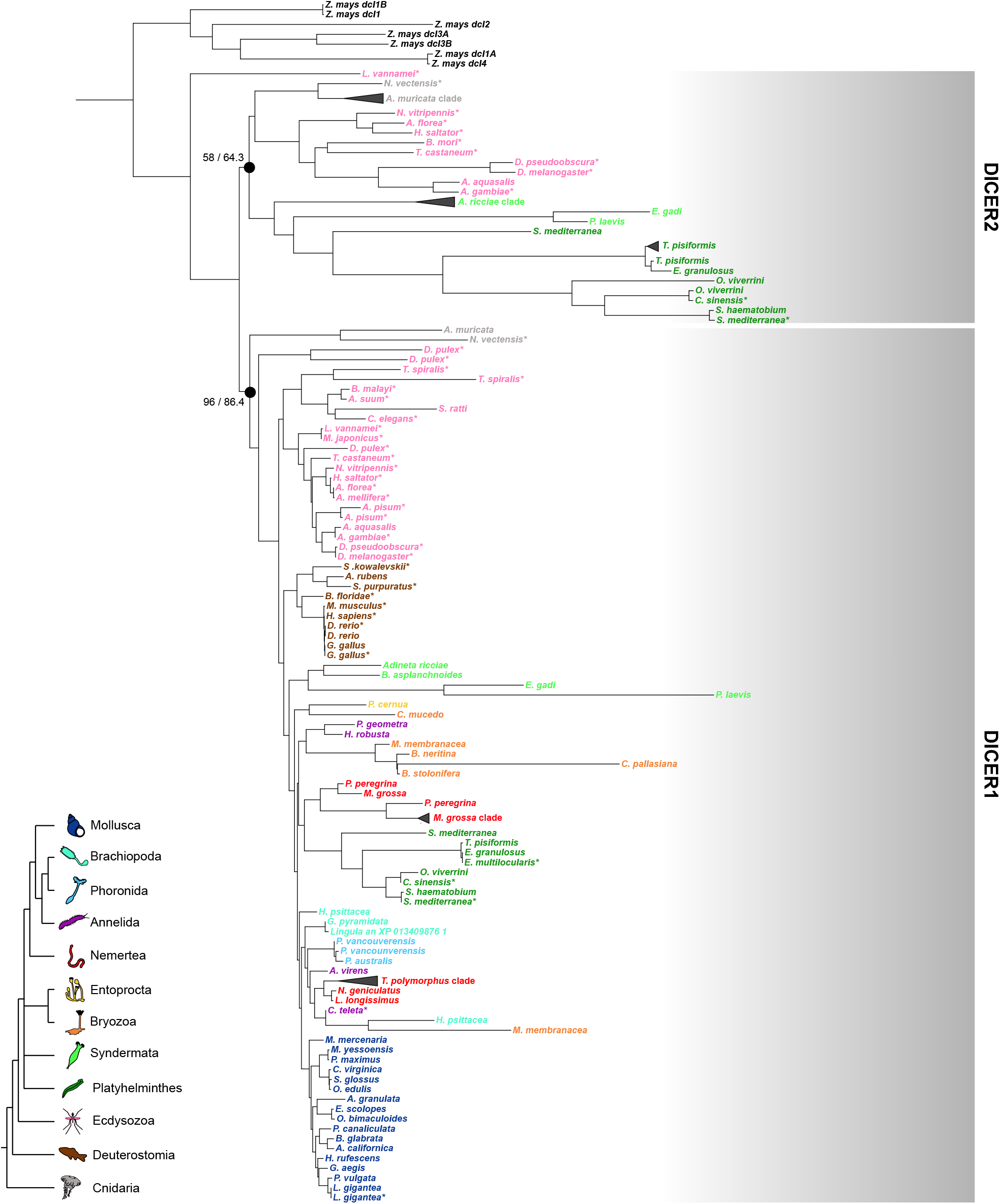
ML tree of the lophotrochozoan DICER proteins. For two marked nodes it is reported respectively the UFBoot and the SH-alrt value. Clades formed by paralogs of the same species were collapsed and represented with a triangle. References species retrieved from the analysis of Mukherjee and colleagues (2013) are marked with an asterisk. Species are colored according to the phylum. The color legend on the bottom left reconstructs the main phylogenetic relationships according to the ML-tree of Kocot and colleagues (2016).

Overall, the DICER family phylogenetic analysis confirmed that the absence of siAGO proteins coincides with the absence of DICER2, and *vice versa*, exception made for Nemertea, Bryozoa and Entoprocta, where we annotated siAGO proteins for most species; however, none of these species featured DICER2. DICER1 and miAGO proteins showed lower evolutionary rates than their paralogous counterparts: the root-to-tip distances of miAGO branches were significantly lower than the root-to-tip distances of siAGO, PIWI1 and PIWI2 branches, and the root-to-tip distances of DICER1 branches were significantly lower than their DICER2 counterparts (supplementary figure S3). We also estimated the ratio of non-synonymous to synonymous substitution rates (*ω*) along the AGO family tree and the DICER family tree: we confirmed that, during the evolution of DICER and AGO family, purifying selection on DICER1 (LRT=34.34, p-value = 4.62 × 10^−9^; supplementary figure S4B) and miAGO (LRT=204,64, p-value = 0; supplementary figure S4A) has intensified compared to the rest of the family tree (Wertheim et al. 2015).

In both phylogenetic analyses the presence of a protein in some phyla was not confirmed by all the species. To understand whether it might be related to the quality of the data, we evaluated the completeness of proteomes, genomes, and transcriptomes: Argonaute and DICER proteins were missing more commonly in transcriptomes than in proteomes or genome assemblies, regardless of completeness (supplementary table S1). All proteomes showed a similar presence/absence pattern and high completeness values, while the completeness of the genomes varied significantly between species. In general, in more complete genome assemblies we were also able to annotate more proteins (supplementary table S1). Overall, phyla that are generally more represented in protein databases (i.e., Mollusca, Annelida, Platyhelminthes) showed a more constant presence/absence pattern between species than under-represented phyla (i.e., Nemertea, Bryozoa, Syndermata). It is worth noting that the annotation of Argonaute and DICER was necessarily dependent on databases of orthologous genes (see Materials and methods).

According to the phylogenomic analysis, some lophotrochozoan phyla lack pivotal proteins related to the endo-siRNA pathway. DICER2 generally processes double stranded RNAs producing two 21 bases siRNA duplexes that overlaps of 19 bases. Similarly, piRNAs produced by the ping-pong cycle go in pairs that overlaps of 10 bases (Antoniewski 2014; Khanal et al. 2022). Thus, siRNAs and piRNAs have a unique signature that can be spot in sncRNA libraries by looking for overlapping pairs of reads. We retrieved eight sncRNA libraries from lophotrochozoan and non-lophotrochozoan species (i.e., *Danio rerio, Apostichopus japonicus, Acropora muricata, Anopheles gambiae, Drosophila melanogaster, Schmitdea mediterranea, Schistosoma japonicum, Crassostrea gigas*) and we sequenced the sncRNA pool of *Notospermus geniculatus*, since no nemertean sncRNA library was available. Using the overlapping_reads.py script (Antoniewski 2014) we calculated the number of read pairs that overlapped for the same number of bases, from 4 to 20 bases. Then, we calculated the Z-Score among the number of read pairs for each overlap group. A Z-score equal to 1 means that the number of reads pairs in that overlap group is one standard deviation higher from the mean size of all the overlap groups of a given species. Taking into consideration only 21 bp reads (i.e., the expected length of endo-siRNAs), Arthropoda (*D. melanogaster* and *A. gambiae*), Cnidaria (*A. muricata*) and Platyhelminthes (*S. mediterranea* and *S. japonicum*) representatives reported a Z-score higher than 1 in the 19-overlap group, corresponding to the siRNA signature (Fig. 3). On the other hand, Deuterostomia (*D. rerio* and *Apostichopus japonicus*), Mollusca (*C. gigas*) and Nemertea (*N. geniculatus*) representatives showed a Z-score lower than 1. Some species, namely *N. geniculatus, A. japonicus, C. gigas*, and *S. mediterranea*, also reported a sharp increase of the Z-score for the 10-overlap group for 21-bp long reads only. A 10-bases overlap would correspond to the piRNA signature, although the length of piRNAs in Arthropoda ranges from 25 to 30 pb. Overall, considering the phylogenomic analysis, all the species that exhibited a complete siRNA pathway (using the presence of DICER2 and siAGO as a proxy) showed a high Z-score in the 19-overlap group (Fig. 3).

**Figure 3.**
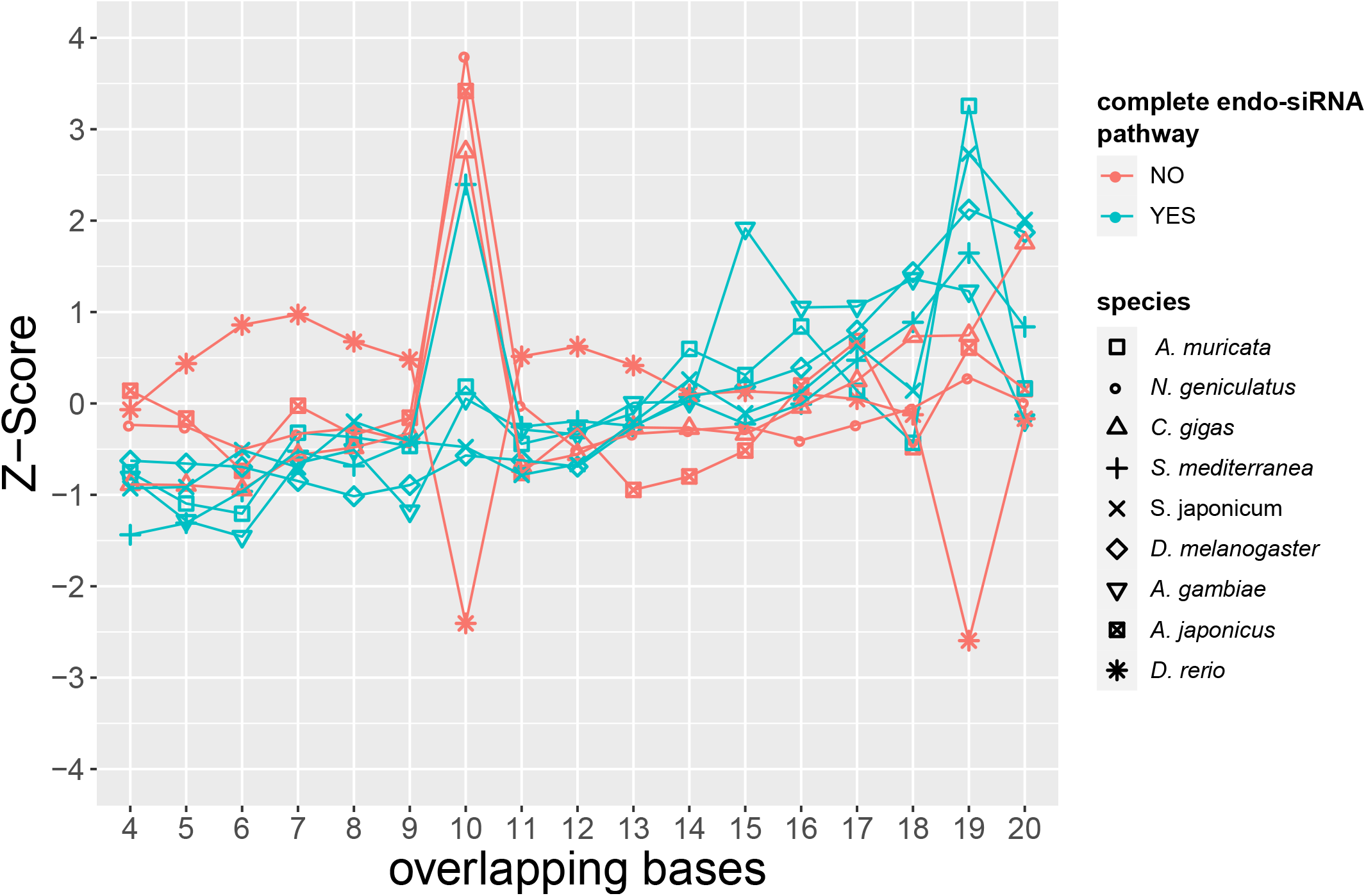
Evaluating the siRNA signature in sncRNA libraries. The plot reports the Z-score between the number of pairs of all possible overlaps. A Z-score greater than 1 means that pairs overlapping of that length are at least a standard deviation more numerous than the mean of all the overlaps. The two colors distinguish species equipped with both endo-siRNA proteins (DICER2 and siAGO) from species that lack at least one of them.

Finally, we investigated whether the range of action of the three sncRNA types is uniform among the three main Metazoa superclades, choosing tree representative for each one (i.e., *C. gigas* for Lophotrochozoa, *A. gambiae* for Ecdysozoa and *D. rerio* for Deuterostomia). We annotated miRNAs, piRNAs and endo-siRNAs evaluating their expression at different sncRNA lengths (supplementary figure S5A). In terms of reads per million (RPM), in all three species miRNAs are the most expressed and their length ranges from 20 – 25 nc. On the contrary, piRNAs were annotated in the in the 24 – 30 nc length range. In most of the cases, endo-siRNAs were not able to discern the endo-siRNA signal from noise, but *A. gambiae* is the species that showed highest expression levels (supplementary figure S4A). We also compared ovary and somatic tissue sncRNAs libraries. As expected, in all three species piRNAs is the class more expressed in the ovaries. (supplementary figure S5B). Overall, in the three species we did not reported notable differences in terms of sncRNA length or differential expression in somatic/ovarian tissue among piRNA and miRNA types.

## Discussion

RNAi pathways play a central role in many molecular aspects, from mRNA regulation to defense mechanisms, and Argonaute proteins are the RNAi key players in all eukaryotes. The expansion and differentiation of this superfamily have reflected the emergence of different RNAi mechanisms, each represented by a sncRNA class loaded by a specific Argonaute. All the phylogenetic analyses agree to divide the eukaryotic Argonaute superfamily in four main families, namely the *Trypanosoma*-AGO family, the WAGO family, the AGO family, and the PIWI family (Höck and Meister 2008; Garcia Silva et al. 2010; Swarts et al. 2014). Excluding the *Trypanosoma*-AGO family, all the other families are represented within Metazoa; moreover, the PIWI and the WAGO family are restricted to animals or even nematodes, respectively (Swarts et al. 2014). It is still uncertain how the four families emerged during eukaryotes evolution. For instance, several eukaryotic clades have a miRNA-like pathway, but it is not clear whether these pathways are analogous or homologous to the metazoan miRNA pathway (Moran et al. 2017). It is likely that RNAi systems diverged from an ancestral siRNA system and, considering the distribution of Eukaryota clades in the four families (Swarts et al. 2014), the divergence took place at least 1.5 billions of years ago (Strassert et al. 2021).

The inferred Argonaute phylogenetic tree confirmed and highly supported the three metazoan Argonaute families (Fig. 1). Moreover, we could identify two distinct groups in the AGO and PIWI, where only one of the two groups is monophyletic. This pattern is confirmed in other Argonaute phylogenetic analyses (Swarts et al. 2014; Praher et al. 2017; Wynant et al. 2017). Recalling the deep divergence of these proteins, the signal might be saturated. Accordingly, most of the nodes at the base of the family are not strongly supported (supplementary figure S1). The same pattern has been observed in the DICER phylogeny, with DICER2 proteins grouped in a polyphyletic group while DICER1 proteins clustered in a well-supported monophyletic clade (Fig. 2) (Mukherjee et al. 2013a). Concordantly, DICER2 and DICER1 are related respectively to siAGO and miAGO proteins. Overall, within each clade the phylogenetic reconstruction is substantially in agreement with the state-of-art animal phylogeny, recalling that the signal has been inferred from single markers.

Almost all lophotrochozoan showed two distant related PIWI proteins (i.e., one AUB-like and one AGO3-like; Fig. 1; supplementary table S1), with the only exception of Neodermata (Platyhelminthes), which lacks the whole piRNA pathway (Fontenla et al. 2021). It is likely that the ping-pong cycle, which has been already described in some mollusks (Jehn et al. 2018), has been maintained in most of lophotrochozoan. The piRNA expression of *C. gigas* is in line with that of *D. rerio* and *A. gambiae* and the differential expression analysis confirms that piRNAs are more expressed in the gonads in all the three clades.

The miRNA pathway appears the most ubiquitous RNAi pathway among Metazoa, almost all species reported a miAGO and a DICER1 protein. For these proteins we even detected lower root-to-tip distances and a decrease of *ω* along their branches, which confirms a higher selective pressure. Therefore, the high conservation of proteins involved in the miRNA pathway reflects the well-known conservation of miRNAs among Metazoa (Tarver et al. 2013). Even in our case, we confirmed that the annotated miRNAs showed similar features in the three reference species; they are 20-25 nt long, they are not more expressed in the ovaries compared to somatic tissue and overall they are by far the most expressed sncRNA class.

Conversely, the endo-siRNA pathway is by far less studied than the miRNA and piRNA ones, which are also more ubiquitous in Metazoa. The absence of siAGO proteins have been reported in Vertebrata (Wynant et al. 2017) and in the annelid *Capitella teleta* (Khanal et al. 2022), where the absence of a siRNA signature in sncRNA libraries have been also confirmed. Our analysis on Argonaute and DICER proteins revealed a unique pattern in the evolution of this pathway. DICER2 and siAGO proteins were retrieved in Platyhelminthes and Syndermata. The two proteins were completely absent in Mollusca, Annelida, Phoronida and Brachiopoda. A unique condition was recovered in Nemertea, Bryozoa and Entoprocta; they are equipped with siAGO proteins (albeit not every species), but DICER2 proteins were not annotated in any of those. Thus, we speculate that Nematoda, Bryozoa and Entoprocta are an intermediate state during the loss of the endo-siRNA pathway in Lophotrochozoa. The phylogenetic relationships between lophotrochozoan phyla are not completely resolved yet. However, a phylotranscriptomic analysis has shown that the best ML tree, inferred by reducing the branch-length heterogeneity, places Nemertea in sister relationship with Mollusca + Phoronida + Annelida + Brachiopoda (Kocot et al. 2016), the group of phyla totally lacking the endo-siRNA pathway. The same tree places Bryozoa and Entoprocta as the sister group of Trochozoa (Nemertea + Mollusca + Phoronida + Annelida + Brachiopoda) and the Polyzoa hypothesis (i.e., monophyly of Bryozoa+Entoprocta) has been recently supported (Khalturin et al. 2022). Therefore, the characterization of endo-siRNA pathway in Nemertea is of paramount importance to achieve a complete figure of the evolution of this pathway. Overall, although the relationships between lophotrochozoan phyla are still uncertain, in Nemertea and Polyzoa the presence of siAGO and absence of DICER2 proteins may have a phylogenetic meaning. We hypothesize that the loss of DICER2 took place at the base of Polyzoa+Trochozoa; subsequently, nemerteans and Polyzoa retained siAGO proteins that were left in the last common ancestor of remaining Trochozoa (Fig. 4). This hypothesis would confirm the phylogenetic position of Polyzoa as sisters of Trochozoa, which is inferred but not highly supported in the latest phylogenetic analyses (Kocot et al. 2016).

**Figure 4.**
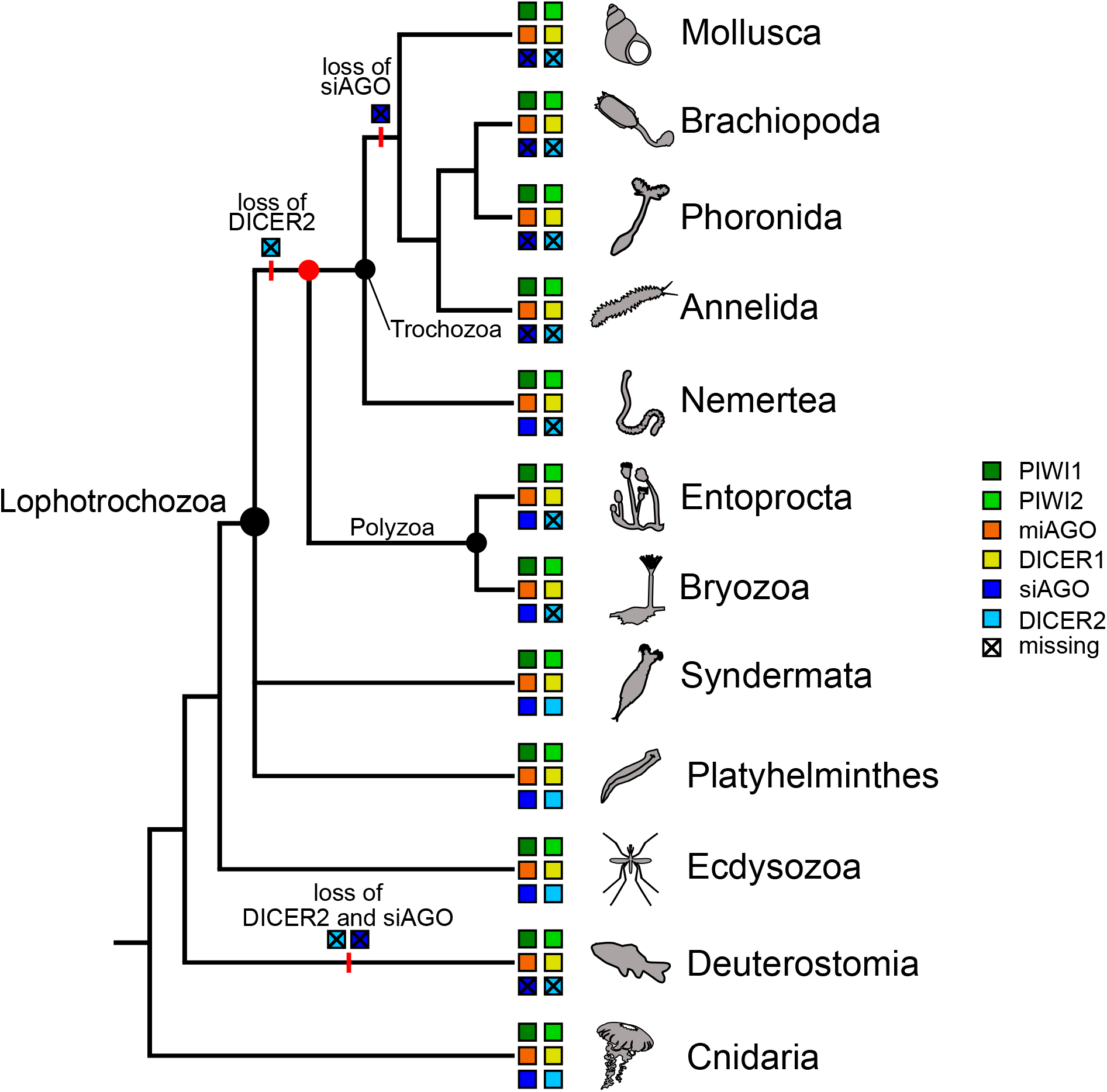
The loss of DICER2 and siAGO along the Metazoa evolution. The lophotrochozoan phylogenetic tree is reconstructed according to the latest Lophotrochozoa phylogenetic analyses (Kocot et al. 2016; Khalturin et al. 2022). The position of Polyzoa as sister group of Trochozoa is weakly supported in the literature (red node), but it would be confirmed by the presence of siAGO and absence of DICER2. Colored boxes represent the presence of the related protein in that clade, while crossed boxes represent the absence of the related protein. Each protein is associated with a color, as reported in the legend.

Indeed, species lacking DICER2 showed less sncRNA pairs overlapping for 19 bp, which is the siRNA signature, compared to species with a complete endo-siRNA pathway. The absence of a siRNA signature in the nemertean *N. geniculatus* is in line with our scenario since the typical signature is due to the DICER2 cleavage on double-stranded RNAs. To our knowledge, an uncomplete endo-siRNA pathway has never been described, at least among Metazoa. Thus, it is difficult to say whether in Nemertea and Polyzoa the siAGO proteins have been coopted to load a different sncRNA type or they have lost their function in the RNAi pathway after the loss of DICER2. Although the miRNA pathway is highly conserved among Metazoa, many other non-canonical pathways have been discovered, which bypass the Microprocessor or the DICER1 step, but are still loaded by miAGOs (Havens et al. 2012). In the same way, nemerteans and polyzoans siAGO proteins might be involved in pathways that do not require the action of DICER2. Alternatively, DICER2 might have been coopted for completely different functions.

Concluding, the loss of siAGO proteins and, more in general, of the endo-siRNA pathway possibly in all Trochozoa have been strongly supported by the joint phylogenetic analysis and analysis of sncRNA libraries. The situation is comparable with that obtained from Deuterostomia, where the absence of the canonical (i.e., involving both DICER and siAGOs) endo-siRNA pathway have already been reported using the lack of annotated siAGO proteins as a proxy (Wynant et al. 2017). Our analysis support this hypothesis, since neither siAGO nor DICER2 homologs were retrieved in deuterostomes (Fig. 1,2). The distribution of the endo-siRNA pathway in Cnidaria (Calcino et al. 2018; Fridrich et al. 2020), Arthropoda and basal Lophotrochozoa (i.e., Platyhelminthes and Syndermata; Fig. 4) suggests that most trochozoans and deuterostomes lost it secondarily. That said, in mammals it has been demonstrated that endogenous dsRNAs can still be processed and included in the RNAi regulation by proteins of the miRNA pathway (Babiarz et al. 2008; Watanabe et al. 2008; Laura et al. 2020). Thus, a different endo-siRNA pathway seems to have evolved from other pathways. In the same way, it is possible that in most Trochozoa endo-siRNAs might not be completely absent, rather processed by different proteins: however, due to the lack of DICER2, they would not display the typical siRNA signature (Park et al. 2021). As for mammals, immunoprecipitation or knockout experiments in Trochozoa might elucidate whether Argonaute proteins can load other sncRNAs types beside piRNAs and miRNAs.

Through the present work we were able to depict the evolution of the main RNAi mechanisms in Metazoa. Unique patterns emerged, such as the uncomplete endo-siRNA pathway in Nemertea and Polyzoa. More efforts should be placed in studying RNAi in less considered phyla, in order to better characterize understudied patterns, casting light on the diversity of RNAi across Metazoa. Comparing different RNAi pathways allowed us to highlight the evolutionary constraints on the miRNA pathway among Metazoa. The conservation and ubiquity of this pathway is likely due to its importance in many regulative systems, and further research should focus on detecting the key features that made the miRNA pathway the main RNAi system.

## Materials and Methods

### Annotation of Argonaute and DICER proteins

Argonaute and DICER proteins were annotated for a wide range of omics-data. We first analyzed all the lophotrochozoan proteomes which were annotated following the NCBI Eukaryotic Genome Annotation Pipeline(Thibaud-Nissen et al. 2016). To increase the sampling in underrepresented clades, we selected 17 assemblies (Table S1) and we predicted gene models with the BRAKER2 automated pipeline (Stanke et al. 2008; Hoff et al. 2019; Brůna et al. 2021). To improve the DICER and Argonaute model prediction, the Metazoa OrthoDB database provided by BRAKER2 was enriched of Argonaute and DICER sequences annotated from the lophotrochozoan NCBI proteomes. We collapsed all the isoforms, keeping the longer ones, using the perl script agat_sp_keep_longest_isoform.pl (Dainat). Lastly, we searched Argonaute and DICER proteins in transcriptomes when phyla showed few assemblies (Table S1). We trimmed the reads with Trimmomatic-0.39 (Bolger et al. 2014) with the following settings: ILLUMINACLIP: TruSeq3-PE.fa:2:30:10 LEADING:3 TRAILING:3 SLIDINGWINDOW:4:15 MINLEN:75; we assembled the transcriptome with Trinity v2.1.1 with default settings (Grabherr et al. 2011); we filtered out contaminants locally aligning the transcripts against the non-redundant protein database (Sayers et al. 2022) with DIAMOND blastp (Camacho et al. 2009; Buchfink et al. 2015) and discarding all the transcripts with a non-metazoan best hit. Coding regions were predicted with Transdecoder v5.5.0 (https://github.com/TransDecoder/TransDecoder), scanning all ORFs for homology using DIAMOND blastp and HMMER (Mistry et al. 2013). Overall, we obtained the coding sequences of 42 lophotrochozoan species and six other metazoan species. Coding sequences were translated into amino acid sequences; then Argonaute and DICER proteins were annotated as follow. We looked for Argonaute proteins by annotating the conserved PIWI and PAZ domains in all 49 species. Domain alignments were retrieved from Pfam (PIWI accession: pfam02171; PAZ accession: pfam02170) (Mistry et al. 2021). Using HMMER, we built a profile for each multiple sequence alignment and we searched the profiles against each (--e-value 10e-6). Only proteins with both domains annotated were considered Argonaute proteins and kept for the phylogenetic analysis.

To annotate DICER proteins we aligned the amino acid sequences against the bilaterian annotated set of Mukherjee and colleagues (Mukherjee et al. 2013b) using blastp. In a second round of filtering, we retrieved the ribonucleases 3 domain alignment from Pfam (accession: pf14622.9), which is the only domain shared between all the metazoan DICERs (Mukherjee et al. 2013b); we searched with HMMER (--e-value 10xe-6) and selected only sequences containing the ribonucleases 3 domain. However, orthologs of the endoribonuclease DROSHA were possibly included among the annotated DICER at this stage. Therefore, we downloaded the DROSHA orthologs from OrthoDB (reference: 9211at3208) (Zdobnov et al. 2021) and we built a custom dataset with both DICER from Mukherjee and colleagues (2013) and DROSHA sequences. We locally aligned the set of putatively annotated DICER proteins against this dataset. We retained proteins whose best five hits were all with DICER orthologs, whereas we discarded proteins with only DROSHA orthologs among the best five hits. No ambiguous results (i.e., proteins showing both DICER and DROSHA within the best five hits) were obtained.

The Argonaute and DICER phylogenetic trees were inferred from datasets being comprised of all the annotated sequences from proteomes, genomes and transcriptomes, with the addition of reference sequences chosen from SwissProt (Fig. 1) or the bilaterian annotated set for the DICER dataset. Datasets were aligned with MAFFT v7.508 (Katoh and Standley 2013), using the options --maxiterate 1000 --localpair. Uninformative columns were masked from the alignments using Gblocks (Castresana 2000), setting -b2= (3 × number of sequences)/5 -b3=10 -b4=5 -b5=a. The ML trees were inferred with IQ-Tree(Nguyen et al. 2015) with the predefined protein mixture model LG+C20+R4. To assess the robustness of the clades we calculated the ultrafast bootstrap approximation (UFBoot) with 1,000 bootstrap replicates (Hoang et al. 2018) and the SH-like approximate likelihood ratio test with 1,000 replicates (Guindon et al. 2010).

We tested whether DICER1 and miAGO proteins have experienced an intensified selection with HyPhy RELAX (Wertheim et al. 2015). We tagged all the branches belonging to DICER1 and miAGO clades as foreground. All the other branches belonging to DICER or AGO family clades were tagged as background. For some proteins we were not able to retrieve the respective coding sequence, namely the *Saccostrea glomerata* miAGO and DICER1, *Anopheles gambiae* DICER2, *Trobolium castaneum* and *Brugia malayi* DICER1. Those proteins were removed during the selection analysis.

The completeness of proteomes, assembly and transcriptomes was evaluated with BUSCO v. 5.4.3 (Simão et al. 2015) using the Metazoa dataset.

### *Notospermus geniculatus* sncRNA libraries sequencing and analysis of sncRNA libraries

Six specimens (three males and three females) of the nemertean Notospermus geniculatus were sampled in June 2018 near Ushimado (Okayama prefecture, Japan). Animals were left in seawater and 7% MgCl_2_·H_2_O (1:1 ratio) for 15’; gonads were then dissected in 7% MgCl_2_·H_2_O on ice and stored in RNAlater (Thermo Fisher Scientific Inc., Waltham, USA), following manufacturer’s instructions. Total RNA was extracted using a standard chloforom:TRI Reagent® (Merck KGaA, Darmstadt, Germany) protocol, following manufacturer’s instructions. The TruSeq Small RNA library kit (Illumina, San Diego, USA) was used to prepare six sRNA libraries that were sequenced on an Illumina HiSeq2500 platform. Both library preparation and high-throughput sequencing were carried out at the Macrogen Inc. facility (Seoul, South Korea).

For this study we sequenced the sncRNA pool from six samples of *N. geniculatus*. These libraries were analyzed along with publicly available sncRNA libraries of other five species. The libraries were selected and downloaded from the Sequence Reads Archive (SRA) provided by NCBI (Table S2). When more samples from the same project were available, libraries were pulled together, obtaining a single fastq file for each species. Adapters and low-quality bases were removed from reads with Cutadapt v3.9.7 (Martin 2011), setting the options -e 0.2 -O 5 --quality-cutoff 6 --discard-untrimmed. Trimmed reads were mapped on the reference genome with Bowtie (Langmead et al. 2009), allowing up to 100 multiple alignments (-m 100). We estimated the distribution of the overlaps between reads with the python script *overlapping_reads*.*py* (https://github.com/ARTbio/tools-artbio/blob/master/tools/small_rna_signatures/overlapping_reads.py; Antoniewski 2014).

For *Crassotrea gigas, Danio rerio* and *Anopheles gambiae* we annotated miRNAs, siRNAs and piRNAs. The tool miRDeep2 (Friedländer et al. 2012) was used to predict miRNAs providing the already annotated miRNA set of that species from MiRGeneDB (Fromm et al. 2022) or miRBase (Kozomara et al. 2019) and the annotated miRNAs of up to five closely related species. The novel predicted miRNAs were reconsidered following the rules established by Fromm and colleagues (2015) and discarding novel miRNAs with a STAR sequence coverage lower than 5 reads. Putative siRNA and piRNA pairs were predicted according to reads overlaps (i.e. siRNA pairs overlap of the read length – 2, piRNA pairs overlap of 10 nucleotides) calculated with *overlapping_reads*.*py*. Pairs of siRNAs and piRNAs were discarded when: one of the paired sRNAs had a coverage lower than 5; the logarithmic ratio of the pair was above 1.5; the pair mapped in a miRNA region. The differential expression of sncRNAs between somatic tissues and ovaries was tested with edgeR (Robinson et al. 2010) using a generalized linear model and a quasi-likelihood F-test (Lund et al. 2012). We chose a strict P-Value threshold of 0.001 to consider a sRNA to be significantly differentially expressed.

## Supporting information

Supplementary information cited through the paper

## Data availability

Raw reads were uploaded to Sequence Read Archive (SRA) under the BioProject PRJNA942081.

## Authors contributions

A.F., F.P. and M.P. designed the research; M.H. and T.S. sampled and prepared the specimens; A.F. and G.C. analyzed the data and wrote the first draft of the article; F.P. and M.P. supervised the project. All authors accepted and contributed to the final version of the article.

## Acknowledgements

This work was supported by the Canziani Bequest and by the Italian Ministry of University and Research PRIN 2020 (2020BE2BC3), both funded to M.P.

