## Supplementary information cited through the paper for "The evolution of RNA interference among Metazoa"

**Supplementary Tables** pag. 2

**Supplementary Figures** pag. 5

| Phylum | Species | PIWI1 | PIWI2 | miAGO | siAGO | DICER1 | DICER2 | Data | Comp. |
|---|---|---|---|---|---|---|---|---|---|
| Mollusca | *Haliotis rufescens* | ✓ | ✓ | ✓ | ✗ | ✓ | ✗ | NCBI Ref. | 99.4% |
| | *Mizuhopecten yessoensis* | ✓ | ✓ | ✓ | ✗ | ✓ | ✗ | NCBI Ref. | 98.6% |
| | *Gigantopelta aegis* | ✓ | ✓ | ✓ | ✗ | ✓ | ✗ | NCBI Ref. | 98.5% |
| | *Pecten maximus* | ✓ | ✓ | ✓ | ✗ | ✓ | ✗ | NCBI Ref. | 98.5% |
| | *Pomacea canaliculata* | ✓ | ✓ | ✓ | ✗ | ✓ | ✗ | NCBI Ref. | 98.2% |
| | *Crassostrea virginica* | ✓ | ✓ | ✓ | ✗ | ✓ | ✗ | NCBI Ref. | 98.1% |
| | *Aplysia californica* | ✓ | ✓ | ✓ | ✗ | ✓ | ✗ | NCBI Ref. | 97.8% |
| | *Ostrea edulis* | ✓ | ✓ | ✓ | ✗ | ✓ | ✗ | NCBI Ref. | 96.8% |
| | *Lottia gigantea* | ✓ | ✓ | ✓ | ✗ | ✓ | ✗ | NCBI Ref. | 96.5% |
| | *Mercenaria mercenaria* | ✓ | ✓ | ✓ | ✗ | ✓ | ✗ | NCBI Ref. | 95.4% |
| | *Octopus bimaculoides* | ✓ | ✓ | ✓ | ✗ | ✓ | ✗ | NCBI Ref. | 94.6% |
| | *Patella vulgata* | ✓ | ✓ | ✓ | ✗ | ✓ | ✗ | NCBI Ref. | 90.5% |
| | *Biomphalaria glabrata* | ✓ | ✓ | ✓ | ✗ | ✓ | ✗ | NCBI Ref. | 88.9% |
| | *Saccostrea glomerata* | ✓ | ✓ | ✓ | ✗ | ✓ | ✗ | NCBI Ref. | 88.9% |
| | *Acanthopleura granulata* | ✓ | ✓ | ✓ | ✗ | ✓ | ✗ | Assemb. | 69.8% |
| | *Euprymna scolopes* | ✓ | ✓ | ✓ | ✗ | ✓ | ✗ | Assemb. | 55.5% |
| Annelida | *Alitta virens* | ✓ | ✓ | ✓ | ✗ | ✓ | ✗ | Assemb. | 69.8% |
| | *Piscicola geometra* | ✓ | ✓ | ✓ | ✗ | ✓ | ✗ | Assemb. | 43.7% |
| | *Helobdella robusta* | ✓ | ✓ | ✓ | ✗ | ✓ | ✗ | NCBI Ref. | 90.2% |
| Brachiopoda | *Lingula anatina* | ✓ | ✓ | ✓ | ✗ | ✓ | ✗ | NCBI Ref. | 98.6% |
| | *Hemithiris psittacea* | ✓ | ✓ | ✓ | ✗ | ✓ | ✗ | Transc. | 90.7% |
| | *Glottidia pyramidata* | ✗ | ✗ | ✗ | ✗ | ✓ | ✗ | Transc. | 75.1% |
| Phoronida | *Phoronis australis* | ✓ | ✓ | ✓ | ✗ | ✓ | ✗ | Assemb. | 68.3% |
| | *Phoronis vancouverensis* | ✓ | ✓ | ✗ | ✗ | ✓ | ✗ | Transc. | 84.9% |
| Nemertea | *Notospermus geniculatus* | ✓ | ✓ | ✓ | ✓ | ✓ | ✗ | Assemb. | 68.4% |
| | *Lineus longissimus* | ✓ | ✓ | ✓ | ✗ | ✓ | ✗ | Assemb. | 69.9% |
| | *Tubulanus polymorphus* | ✓ | ✗ | ✗ | ✓ | ✓ | ✗ | Transc. | 71.8% |
| | *Malacobdella grossa* | ✓ | ✓ | ✗ | ✓ | ✓ | ✗ | Transc. | 91.7% |
| | *Paranemertes peregrina* | ✗ | ✗ | ✗ | ✓ | ✓ | ✗ | Transc. | 80.5% |
| Bryozoa | *Cryptosula pallasiana* | ✓ | ✓ | ✓ | ✓ | ✓ | ✗ | Assemb. | 83.5% |
| | *Cristatella mucedo* | ✗ | ✓ | ✗ | ✓ | ✓ | ✗ | Assemb. | 62.1% |
| | *Membranipora membranacea* | ✓ | ✓ | ✓ | ✗ | ✓ | ✗ | Assemb. | 54.5% |
| | *Bugulina stolonifera* | ✓ | ✗ | ✓ | ✗ | ✓ | ✗ | Assemb. | 54.0% |
| | *Bugula neritina* | ✓ | ✗ | ✓ | ✗ | ✓ | ✗ | Assemb. | 50.7% |
| Entoprocta | *Pedicellina cernua* | ✓ | ✓ | ✓ | ✓ | ✓ | ✗ | Transc. | 90.4% |
| Syndermata | *Adineta ricciae* | ✓ | ✓ | ✓ | ✓ | ✓ | ✓ | Assemb. | 79.6% |
| | *Pomphorhynchus laevis* | ✓ | ✓ | ✗ | ✗ | ✓ | ✓ | Assemb. | 43.2% |
| | *Brachionus asplanchnoidis* | ✓ | ✓ | ✓ | ✓ | ✓ | ✗ | Assemb. | 85.4% |
| | *Echinorhynchus gadi* | ✓ | ✓ | ✓ | ✓ | ✓ | ✓ | Transc. | 58.6% |
| Platyhelminthes | *Echinococcus granulosus* | ✗ | ✗ | ✓ | ✓ | ✓ | ✓ | NCBI Ref. | 69.2% |
| | *Schistosoma haematobium* | ✗ | ✗ | ✓ | ✓ | ✓ | ✓ | NCBI Ref. | 77.1% |
| | *Opisthorchis viverrini* | ✗ | ✗ | ✓ | ✓ | ✓ | ✓ | NCBI Ref. | 64.7% |
| | *Taenia pisiformis* | ✗ | ✗ | ✓ | ✓ | ✓ | ✓ | Assemb. | 36.3% |
| | *Schmidtea mediterranea* | ✓ | ✓ | ✓ | ✓ | ✓ | ✓ | Assemb. | 46.5% |
| others | *Gallus gallus* | ✓ | ✗ | ✓ | ✗ | ✓ | ✗ | NCBI Ref. | 95.3% |
| | *Danio rerio* | ✓ | ✓ | ✓ | ✗ | ✓ | ✗ | NCBI Ref. | 99.6% |
| | *Asterias rubens* | ✓ | ✓ | ✓ | ✗ | ✓ | ✗ | NCBI Ref. | 98.7% |
| | *Anopheles aquasalis* | ✓ | ✓ | ✓ | ✓ | ✓ | ✓ | NCBI Ref. | 99.0% |
| | *Strongyloides ratti* | ✗ | ✗ | ✓ | ✓ | ✓ | ✗ | NCBI Ref. | 69.2% |
| | *Acropora muricata* | ✓ | ✓ | ✓ | ✓ | ✓ | ✓ | Assemb. | 60.4% |

**Table S1**. Presence and absence of Argonaute and DICER proteins. Colored dots refer to colors assigned to the different animal phyla in Figures 1 and 2. For the four Argonaute proteins and the two DICER proteins, a green check marks the species where the protein has been annotated, a red cross marks species where the protein is absent. The “Data” column reports the type of data, proteome annotated following the NCBI

Eukaryotic Genome Annotation Pipeline ("NCBI Ref."), a genome assembly ("Assemb."), or a transcriptome ("Transc."). The last column reports the BUSCO completeness score.

| Species | Data | Accession numbers |
|---|---|---|
| *Haliotis rufescens* | NCBI Ref. | GCF_023055435.1 |
| *Mizuhopecten yessoensis* | NCBI Ref. | GCF_002113885.1 |
| *Gigantopelta aegis* | NCBI Ref. | GCA_016097555.1 |
| *Pecten maximus* | NCBI Ref. | GCF_902652985.1 |
| *Pomacea canaliculata* | NCBI Ref. | GCF_003073045.1 |
| *Crassostrea virginica* | NCBI Ref. | GCF_002022765.2 |
| *Aplysia californica* | NCBI Ref. | GCF_000002075.1 |
| *Ostrea edulis* | NCBI Ref. | GCF_023158985.2 |
| *Lottia gigantea* | NCBI Ref. | GCF_000327385.1 |
| *Mercenaria mercenaria* | NCBI Ref. | GCF_014805675.1 |
| *Octopus bimaculoides* | NCBI Ref. | GCF_001194135.2 |
| *Patella vulgata* | NCBI Ref. | GCF_932274485.1 |
| *Biomphalaria glabrata* | NCBI Ref. | GCF_000457365.1 |
| *Saccostrea glomerata* | NCBI Ref. | GCA_003671525.1 |
| *Acanthopleura granulata* | Assemb. | GCA_016165875.1 |
| *Euprymna scolopes* | Assemb. | GCA_024364805.1 |
| *Alitta virens* | Assemb. | GCA_932294295.1 |
| *Piscicola geometra* | Assemb. | GCA_943735955.1 |
| *Helobdella robusta* | NCBI Ref. | GCF_000326865.1 |
| *Lingula anatina* | NCBI Ref. | GCF_001039355.2 |
| *Hemithiris psittacea* | Transc. | SRX731469 |
| *Phoronis australis* | Assemb. | GCA_002633005.1 |
| *Phoronis vancouverensis* | Transc. | SRX731474 |
| *Notospermus geniculatus* | Assemb. | GCA_002633025.1 |
| *Lineus longissimus* | Assemb. | GCA_910592395.2 |
| *Tubulanus polymorphus* | Transc. | SRR1275406 |
| *Malacobdella grossa* | Transc. | SRX731465 |
| *Paranemertes peregrina* | Transc. | SRX731466 |
| *Cryptosula pallasiana* | Assemb. | GCA_945261195.1 |
| *Cristatella mucedo* | Assemb. | GCA_009760855.2 |
| *Membranipora membranacea* | Assemb. | GCA_914767715.1 |
| *Bugulina stolonifera* | Assemb. | GCA_935421135.1 |
| *Bugula neritina* | Assemb. | GCA_010799875.2 |
| *Pedicellina cernua* | Transc. | https://datadryad.org/stash/dataset/doi:10.5061/dryad.30k4v |
| *Adineta ricciae* | Assemb. | GCA_900240375.1 |
| *Pomphorhynchus laevis* | Assemb. | GCA_012934845.2 |
| *Brachionus asplanchnoidis* | Assemb. | GCA_025491095.1 |
| *Echinorhynchus gadi* | Transc. | SRR2131254 |
| *Echinococcus granulosus* | NCBI Ref. | GCF_000524195.1 |
| *Schistosoma haematobium* | NCBI Ref. | GCF_000699445.3 |
| *Opisthorchis viverrini* | NCBI Ref. | GCF_000715545.1 |
| *Taenia pisifromis* | Assemb. | GCA_023968675.1 |
| *Schmidtea mediterranea* | Assemb. | GCA_022537955.1 |
| *Gallus gallus* | NCBI Ref. | GCF_016699485.2 |

| | | |
|---|---|---|
| *Danio rerio* | NCBI Ref. | GCF_000002035.6 |
| *Asterias rubens* | NCBI Ref. | GCF_902459465.1 |
| *Anopheles aquasalis* | NCBI Ref. | GCF_943734665.1 |
| *Strongyloides ratti* | NCBI Ref. | GCF_001040885.1 |
| *Acropora muricata* | Assemb. | GCA_014634545.1 |

**Table S2.** Accession numbers of the NCBI RefSeq proteomes, genome assemblies and transcriptomes used.

| Species | Tissue | SRA |
|---|---|---|
| *Anopheles gambiae* | ovary | SRR11713497-9 |
| | midgut | SRR11713500-2 |
| *Crassostrea gigas* | ovary | SRR6489633-4 |
| | muscle | SRR6489635-6 |
| *Danio rerio* | ovary | SRR1265755-7 |
| | gut | SRR1265746-8 |
| *Schmidtea mediterranea* | whole animal | SRR13154740-1 |
| *Schistosoma japonicum* | whole animal | SRR2927289-94 |
| *Drosophila melanogaster* | ovary | ERR3721467-9 |
| *Apostichopus japonicus* | body wall | SRR18918495-500 |

**Table S3.** List of species whose sncRNA libraries were analyzed, reporting the SRA codes used and the sampled tissues.

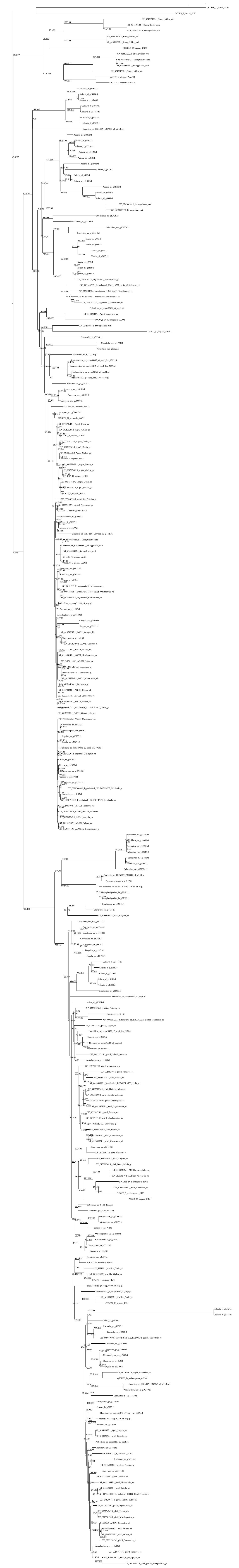

**Figure S1**. ML tree of the lophotrochozoan Argonuaute proteins. For each node it is reported the SH-alrt and UFBoot value

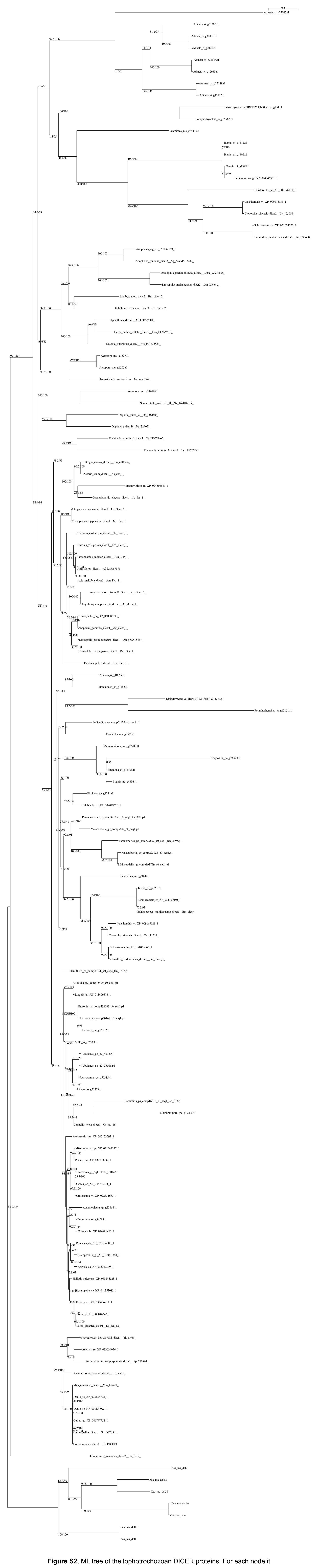

**Figure S2**. ML tree of the lophotrochozoan DICER proteins. For each node it is reported the SH-alrt and UFBoot value

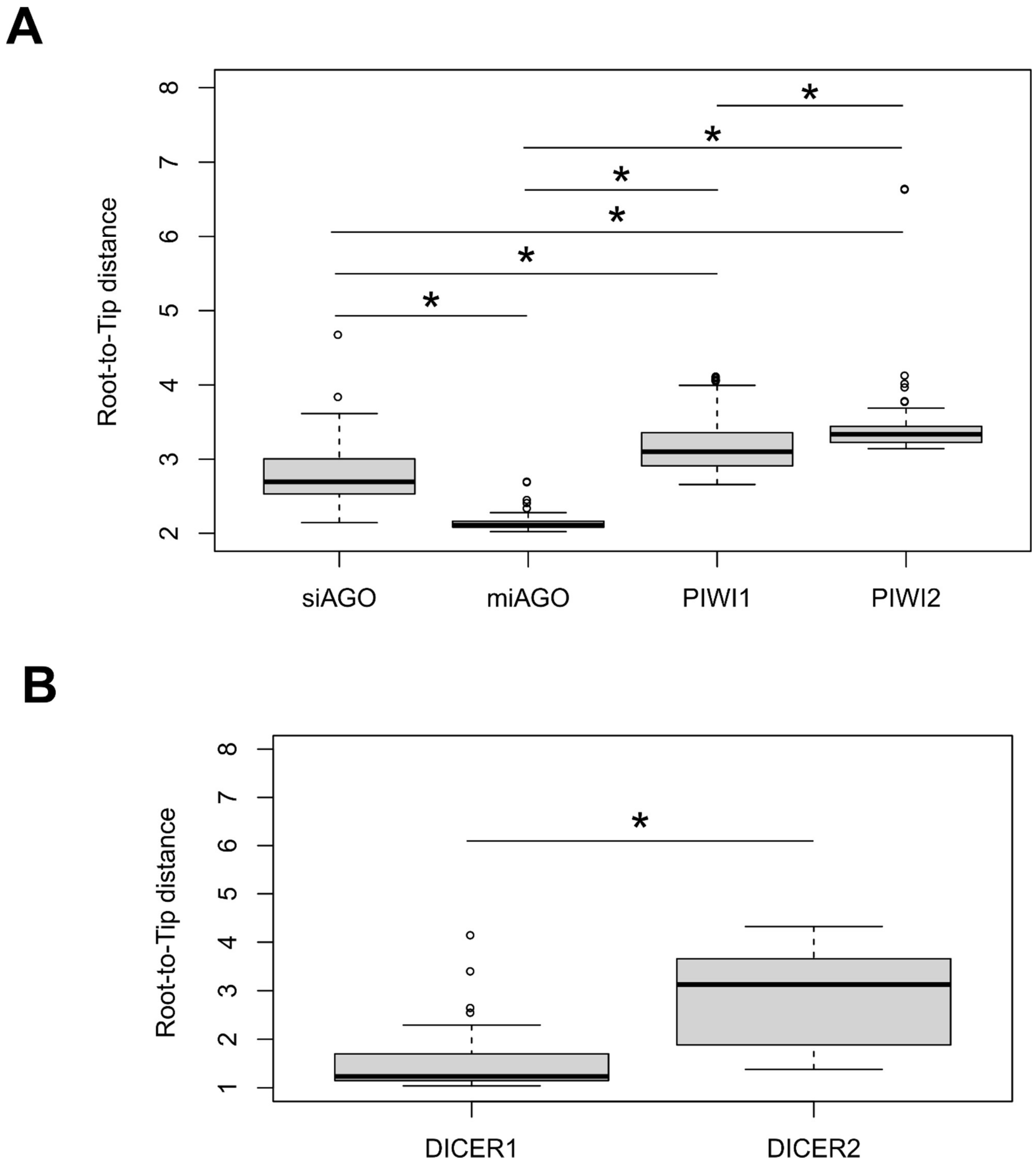


**Figure S3.** Root to tip distance distribution of Argonaute and DICER proteins. The distance of each tip from the root has been calculated by summing all the branch-lengths from the root to the tip. Significance between groups has been evaluated through an ANOVA test, followed by a pairwise t-test. All comparisons resulted significant. (A) Boxplot reporting the root to tip distance distribution of the four Argonaute proteins. Significance between groups was evaluated through an ANOVA test followed by a Tukey's HSD test. All comparisons resulted significant (B) Boxplot reporting the root to tip distance distribution of DICER proteins. According to the Student's t-test, the comparison between the two groups resulted significant.

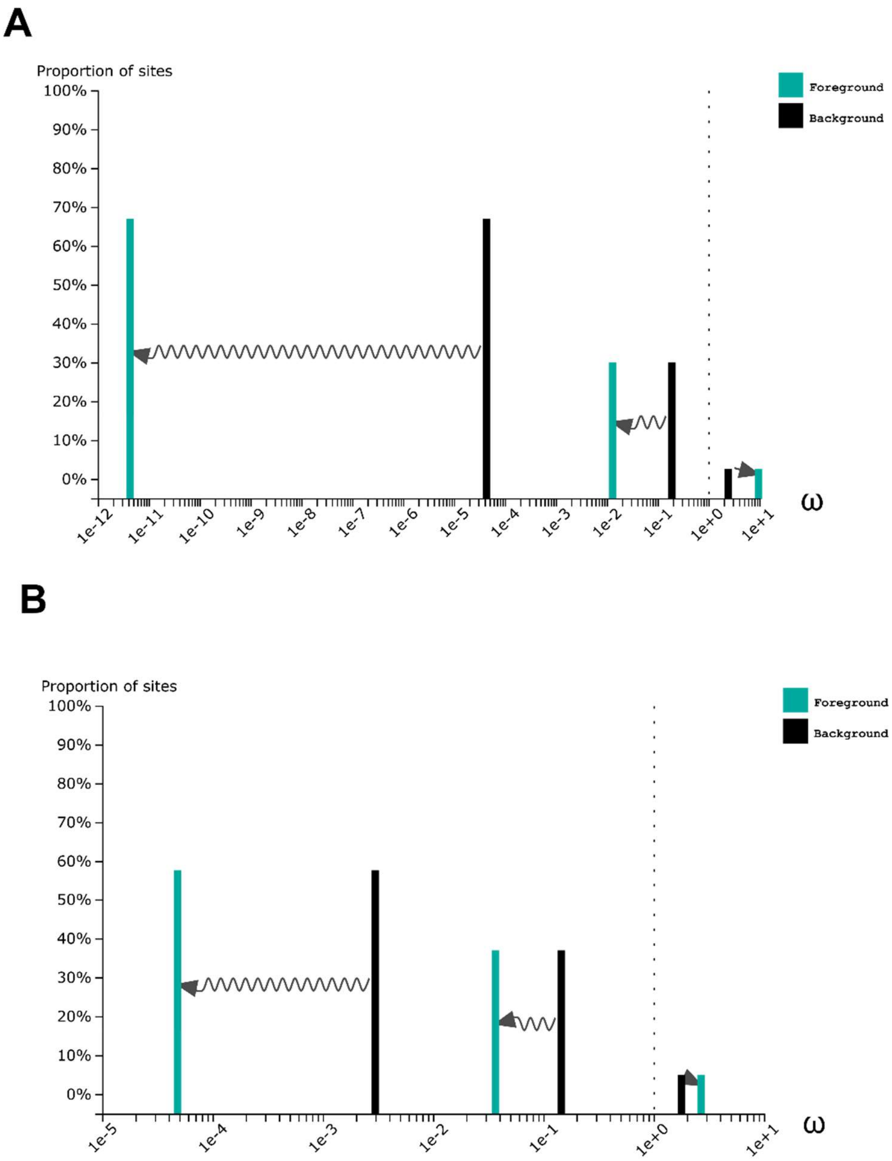


**Fig. S4.** Sites distribution of the three ω classes under the RELAX alternative model. (A) Plot of the RELAX analysis on the AGO family**.** (B) Plot of the RELAX analysis on the DICER family

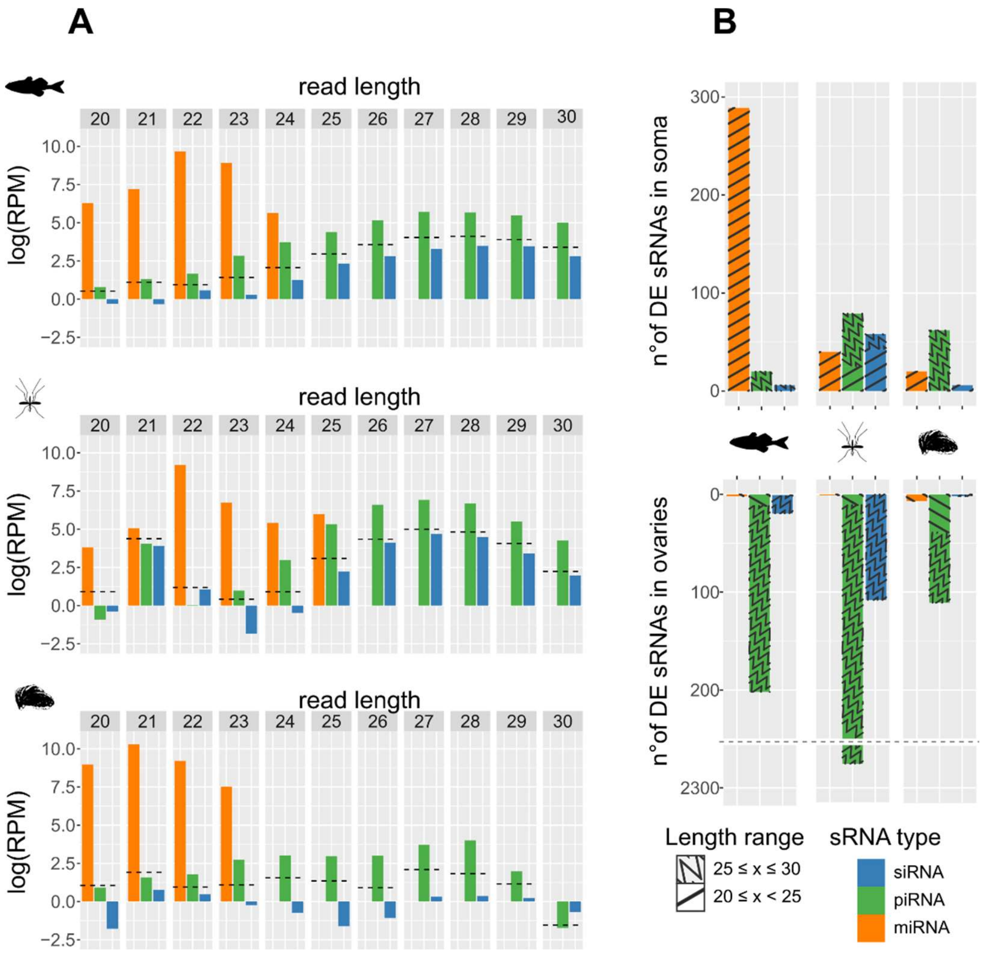


**Figure S5**. Expression of miRNAs (orange), piRNAs (green) and endo-siRNA (blue) at different lengths and in two tissues. (A) Barplots report the expression of the three sRNA types expressed in reads per millions (RMP) on a logarithmic scale. sncRNAs were divided by length. Horizontal lines represent the background signal. To calculated the background signal we excluded reads annotated as miRNAs, siRNAs and piRNAs and computed the highest reads per million value (RPM) among the remaining overlap groups for each read length. We took that value as the background expression level of sRNA for that read length. We expect bona fide miRNAs, piRNAs and siRNAs to be more expressed than the background level in terms of RPM (B) The barplot reports the number of sRNAs significantly more expressed in the ovary against a somatic tissue and vice versa.
